# Holocene North Asian gene flow and cryptic substructure in the genome history of Nilgiri tribes

**DOI:** 10.64898/2026.09.09.750264

**Authors:** Shailesh Desai, Manoj Kumar Tharu, Pratik Pandey, Anjana Welikala, Rakesh Tamang, Gyaneshwer Chaubey

## Abstract

The Nilgiri Hills of the Western Ghats harbour several historically isolated tribal populations whose genomic history remains incompletely resolved. Here, we present a comprehensive analysis of whole-genome and complete mitochondrial data from three major Nilgiri groups i.,e, Toda, Kota, and Kurumba, in the context of a broad Eurasian reference panel. Autosomal analyses reveal previously unrecognised fine-scale substructure, with both Kota and Kurumba resolving into two genetically distinct clusters (Kota1/Kota2 and Kurumba1/Kurumba2). Principal component, ADMIXTURE, and fineSTRUCTURE analyses identify Toda as the most genetically drifted Nilgiri group, whereas Kota2 exhibits elevated Austroasiatic-related ancestry. qpWave analysis indicates that the five genetic clusters require at least three independent ancestry streams, while qpAdm further differentiates the groups based on Onge-, Indus-Periphery-, and Western-Steppe-MLBA-related ancestry. Toda shows the highest Indus-Periphery-related and lowest Onge-related ancestry among the Nilgiri groups. A distinctive ancestry component shared by Kurumba2, Kota1, Gaud and the Sri Lankan Vedda suggests deeper ancestral connections across the southern Indian Ocean region. Mitochondrial genomes reveal strong founder effects and lineage-specific patterns, including R5a2b in Toda, M3a1 in Kota, and multiple M subclades together with the rare C4a2c1 lineage in Kurumba. Notably, C4a2c1, previously understudied in South India, shows phylogenetic and ancient-DNA continuity with North Asian and Siberian lineages. Bayesian dating suggests its arrival in the Indian subcontinent during the early–mid Holocene (∼7 ka), followed by local diversification after ∼5 ka. Together, these findings reveal a complex history of long-range Holocene maternal gene flow and subsequent isolation-driven differentiation, providing new insights into the genomic history of South Indian tribal populations.

## Introduction

The Nilgiri plateau of the Western Ghats is one of the most ethnographically distinctive highland regions of South India. Its indigenous communities, particularly the Toda, Kota and Kurumba, have long attracted anthropological attention because of their small census sizes, endogamous social organisation, specialised subsistence practices and relative geographic isolation (Breeks 1873; Thurston 1909; Ghosh et al., 1977; Vishwanathan et al., 2004). These Nilgiri populations are very interesting due to its complex history, like Kurumba, are divided into many small clans, but they are believed to be descendants of Great Kingdom empire of Pallava, The Toda people exhibit one of the highest frequencies of lactase persistence on the Indian subcontinent, with levels comparable to those found in Northern European populations, despite having a steppe ancestry of only around 7.8% (Kerdoncuff et al., 2026). These groups occupy different social hierarchies, each function complementary to each other and co-existed for a long time (Ghosh et al., 1977). Linguistic and cultural evidence places them within the broader Dravidian-speaking tribal landscape of southern India, yet their precise relationships to one another and to neighbouring populations have remained only partially resolved by earlier genetic surveys that relied on limited autosomal markers or uniparental loci (Cordaux et al. 2003; Vishwanathan et al. 2004).

Recent large-scale whole-genome resources, most notably the GenomeAsia 100K Consortium data, now permit a high-resolution examination of these groups. Previous work has established that the majority of South Asian genetic variation can be modelled as a mixture of three deeply diverged ancestral components such as Ancestral South Indian (AASI/Onge-related), Iranian farmer-related (often approximated by Indus Periphery samples), and Steppe pastoralist-related ancestry with additional East Asian-related contributions in Austroasiatic and Tibeto-Burman speakers (Reich et al. 2009; Narasimhan et al. 2019; GenomeAsia 100K Consortium 2019). Within this framework, South Indian tribal populations typically retain higher proportions of AASI-related ancestry and show signals of strong genetic drift and founder events, consistent with long-term isolation and small effective population sizes. However, nilgiri groups remained understudied despite their very uniqueness.

Here we combine complete mitochondrial genomes, genome-wide SNP data and formal statistical modelling to reconstruct the maternal and autosomal history of the Toda, Kota and Kurumba. We pay particular attention to (i) the unexpected presence of the North Asian/Siberian-associated mitochondrial haplogroup C4a2c1 in the Kurumba, (ii) previously unrecognised internal genetic substructure within the Kota and Kurumba, (iii) quantitative differences in ancestral source proportions among the Nilgiri groups, and (iv) potential haplotype sharing with Sri Lankan indigenous populations. The results demonstrate both deep local continuity and surprising long-distance connections, underscoring the complexity of South Indian tribal genome history.

## Methodology

### mtDNA analysis

To investigate the overall mitochondrial haplogroup composition of the three major Nilgiri tribal populations, complete mitochondrial genomes were extracted from the GenomeAsia GenomeAsia100K Consortium (2019) dataset using the bcftools consensus tool for Toda (N = 20) and Kota (N = 8). For Kurumba, which was represented only in the dataset published by Mait and colleagues, mitochondrial sequences were retrieved from the NCBI database (N = 108) (Kumar et al., 2009) as complete mitochondrial genomes were not available for the Kurumba individuals in the GenomeAsia dataset.

In addition, comparative mitochondrial sequences were collected based on the haplogroup affinities of each population to investigate the geographic distribution, potential lineage origins, and maternal relationships of the Nilgiri populations with other populations. The complete list of comparative sequences is provided in Supplementary Table 1.

### Ancient DNA analysis of mtDNA

During the investigation of mitochondrial haplogroup diversity, the C4a2c haplogroup was identified in the Kurumba population, specifically within the Jenu Kurumba clan. This haplogroup has been reported primarily in populations from Northeast Asia and Siberia. To investigate the possible migration history and maternal origin of the C4a2c lineage in the Kurumba population, we conducted a comprehensive literature survey and used MitoMapper to identify previously reported occurrences of this haplogroup. We additionally searched the AmtDB (Ehler et al., 2019) and Allen Ancient DNA Resource (AADR) databases (Mallick et al., 2024) and identified C4a2c in 13 samples/populations (details are provided in Supplementary Table 1). Among these, only one sample (AADR: I18744) had a complete mitochondrial sequence directly available.

For the remaining samples, mitochondrial genomes were reconstructed through an ancient DNA analysis pipeline. The sample reported by Kumar et al. (2022) was obtained as a BAM file and subsequently processed using schmutzi (Renaud et al., 2015). For the other samples, raw sequencing data were downloaded and processed using AdapterRemoval (Schubert et al., 2016) which was used to remove adapter sequences and low-quality reads. Only collapsed reads were retained for downstream analyses, using a minimum base quality of 20 and a minimum read length of 25 bp.

The retained reads were aligned to the mitochondrial reference genome using BWA aln with the parameters -l 1024, -n 0.01, and -o 2. The resulting BAM files were subsequently processed to remove duplicate reads using the samtools markdup tool. The final BAM files were used with schmutzi to estimate mitochondrial DNA contamination and reconstruct the mitochondrial consensus sequences. Only sequences showing no evidence of contamination were retained for subsequent analyses.

To further assess sequence quality and identify potentially spurious mutations, the reconstructed mitochondrial sequences were manually inspected and curated (Supplementary File 1). The sequences were evaluated in a parsimony framework to identify mutations that were inconsistent with the expected phylogenetic relationships and to assess the overall reliability of each reconstructed sequence (Supplementary File 1).

In order to understand the time split between different lineages, we used Beast2 for Bayesian inference analysis. First all aligned sequences were used for building xml files using beauti, and beast were used to run analysis with 50 Million MCMC chains and with HKY+I+G model and mutation rate of 2.514E-08 (Silva et al., 2017). Resulted log files were checked using Tracer and ensured that each parameter ESS remained more than 200 and final trees were annotated using Treeannottor. For aDNA specifically we used sample date as tip date in Beast2 to infer the demography (Bouckaert et al., 2019).

## Autosomal Analysis

### Autosomal data analysis

For autosomal-level analyses, whole-genome data were retrieved from the GenomeAsia 100K Consortium, which comprises a major representation of South Asian populations. These data were merged with the laboratory dataset, which includes data from the Mait dataset (Metspalu et al., 2010) and the Human Genome Diversity Panel (HGDP) (Bergström et al., 2020). In addition, recently published whole-genome data from the Sinhalese and Vedda populations of Sri Lanka were incorporated (Urban Aragon et al., 2025). The resulting dataset comprised 2,431 individuals representing 167 populations from across the globe. This dataset is hereafter referred to as the Main dataset and was used for the majority of the downstream autosomal analyses. Importantly in all downstream Analysis, we have divided the global population into different Geographical regions, but only the Indian population has been further divided into language wise such as Indo-European, Dravidian, Tibeto-Burman and AustroAsiatic.

### Population Structure and Admixture

To investigate the genetic structure and patterns of genetic variation among the three major Nilgiri tribal populations, Kota, Toda, and Kurumba. We first performed quality control and linkage disequilibrium (LD) pruning of the Main dataset. Data management and quality-control procedures were performed using PLINK2 (Chang et al., 2015). To minimize the influence of background LD on downstream principal component analysis (PCA) and ADMIXTURE analyses (Alexander et al., 2009), we removed one SNP from each pair of variants showing strong LD (r^2^ > 0.4) within a sliding window of 200 SNPs, with the window shifted by 25 SNPs at each step. Following LD pruning, a total of 163,932 high-quality SNPs were retained for subsequent PCA and ADMIXTURE analyses.

Principal component analysis was performed using the smartpca program implemented in the EIGENSOFT package (Price et al., 2006), using default parameters. PCA was used to characterize broad patterns of genetic variation and to assess the relative genetic affinities of the Nilgiri populations with populations from other geographic regions. For comparative interpretation, tribal and caste populations were broadly categorized according to their linguistic affiliations, including Indo-European, Austroasiatic, Trans-Himalayan, and Dravidian language groups. We also divided the Kota and Kurumba populations into two distinct groups within them, as indicated in PCA and later confirmed by Admixture and fineSTRUCTURE (Lawson et al., 2012).

We subsequently performed model-based ancestry estimation using ADMIXTURE (Alexander et al., 2009) on the LD-pruned dataset. For each value of K ranging from 2 to 15, ADMIXTURE was run 10 independent times using different random seeds to assess the consistency of the inferred ancestry components. Based on the overall clustering patterns and model evaluation, K = 11 was considered the most informative clustering level for interpreting the population structure.

Based on the patterns observed in PCA and ADMIXTURE analyses, Kota and Kurumba were subdivided for population-level analyses to minimize their potential influence on estimates of population genetic relationships. To further investigate the genetic affinities of the Nilgiri populations, outgroup f3-statistics (Patterson et al., 2012) were calculated in the form f3(Nilgiri Tribe, X; Yoruba), where X represented each comparative population and Yoruba was used as the outgroup. Higher f3 values were interpreted as greater shared genetic drift between the Nilgiri population and the corresponding comparative population.

### Haplotype-based population structure

To investigate population structure at the haplotype level, genotype data were phased using Beagle v3.3.2 (Browning et al., 2021). The phased dataset was subsequently analysed using fineSTRUCTURE (Lawson et al., 2012), which employs haplotype sharing and a Bayesian clustering framework to identify genetically homogeneous groups and infer fine-scale population structure. The analysis was performed using an MCMC framework with a 10-million-iteration burn-in period, followed by the specified MCMC sampling iterations. The resulting fineSTRUCTURE output was subsequently used to generate a maximum-likelihood (ML) tree, which was constructed using MEGA v7 (Kumar et al., 2016). Moreover co-ancestry matrices were used to understand the haplotype sharing between different groups.

### Founder events and haplotype sharing

To investigate signals of founder events and recent demographic history, we applied ASCEND (Allele Sharing Correlation for Estimation of Nonequilibrium Demography) (Tournebize et al., 2022) to estimate the founder event date and intensity for the Kota1 and Toda populations, as Kota2 and both Kurumba group were having sample size below 5. These estimates were used to assess the timing and relative strength of population contraction or founder-related demographic events.

To further investigate potential historical gene flow and shared ancestry between the Different Nilgiri groups with other populations, we performed IBD analysis. After calculating the IBD, first it was arranged to make a sum and average for population comparison for segment length and segment numbers. IBD was performed to understand the relatedness with other populations.

### Siberian or Northeast Asian Ancestry in Indian Populations

Following the identification of the C4a2c1 mitochondrial haplogroup exclusively in the Kurumba population, and its close maternal affinity with lineages predominantly reported from Siberia and Northeast Asia, we sought to investigate whether the Kurumba, as well as the other Nilgiri tribal populations, possess any distinctive Northeast Asian or Siberian-related ancestry in their autosomal genomes compared with other South Asian populations. To address this question, and to further investigate patterns of shared genetic drift and possible ancestral contributions, we performed qpWave, qpAdm, and outgroup f3-statistics. For these analyses, the Main dataset was merged with selected individuals and populations from the Allen Ancient DNA Resource (AADR), focusing on samples that were particularly relevant to the ancestry and geographic history of the Nilgiri populations. The resulting dataset comprised 2,831 individuals representing 229 populations. The merged PLINK dataset was subsequently converted into EIGENSTRAT format for downstream f-statistic-based analyses. During the conversion using convertf, we encountered a limitation in which the family and individual identifiers in the PLINK fam file were restricted to a maximum of 39 characters. To preserve the complete sample and population identifiers, we developed a custom script that generated a temporary fam file containing shortened identifiers for the conversion step. Following successful conversion to EIGENSTRAT format, these temporary identifiers were remapped to the original identifiers using the original fam file, thereby retaining the complete sample and population names for all downstream analyses.

### qpWave Analysis

To investigate whether the three Nilgiri tribal populations share a common ancestral genetic background or require multiple independent ancestral streams to explain their genetic differentiation, we performed qpWave analysis (Haak et al., 2015). qpWave evaluates the minimum number of ancestry streams connecting the tested populations to a set of outgroup or right populations, thereby allowing assessment of whether genetically differentiated populations can be explained by a common ancestral source or require additional independent ancestry streams. We first tested a simple model in which Toda, Kota, and Kurumba were each treated as a single population, without considering the internal genetic structure identified through PCA and fineSTRUCTURE analyses. We subsequently incorporated the genetic substructure identified in these analyses by dividing the populations into their corresponding genetic clusters, including Kota1, Kota2, Kurumba1, and Kurumba2, and repeated the qpWave analyses to determine whether these subgroups could be explained by the same or distinct ancestral streams. Finally, we incorporated geographically and genetically neighbouring South Asian populations into the analysis to evaluate the position of the Nilgiri populations within the broader regional genetic landscape. These models allowed us to assess the minimum number of independent ancestry streams required to explain the observed genetic relationships among the Nilgiri populations and their regional counterparts.

### qpAdm Analysis

To further investigate the ancestral composition of the Nilgiri populations, we performed qpAdm analyses (Haak et al., 2015) using the ADMIXTOOLS framework. We tested alternative two- and three-source ancestry models to evaluate whether the target populations could be explained as mixtures of selected ancestral populations. The models included Onge, Indus_Periphery_HighCoV, and Western_Steppe_MLBA as candidate sources, representing major ancestral components relevant to South Asian population history (Kerdoncuff et al., 2025). Model support was assessed using the qpAdm P-value, with models having P > 0.01 considered statistically compatible with the data. We compared the inferred ancestry proportions across the Nilgiri populations and selected neighbouring populations to identify similarities and differences in their ancestral composition. These analyses were used to assess whether the Nilgiri populations possess distinctive ancestry components relative to surrounding South Asian populations

### D statistics and F3 Outgroups

First to understand the general affinity of each group of Nilgiri tribe, we performed the outgroup Analysis using (YRI, X, Nilgiri Tribe), whereas to investigate the specific affinity towards the East Asia we used D statistics using D(YRI, East_Asia Population, Paniya, Kurumba1), where significant positive D will mean gene flow to the Kurumba1 and East Asian Populations. However, Paniya already possesses more AASI related component which is more closely related to East Asian lineage, thus to avoid bias we also tested other several combinations specifically selecting population from the North part of India and also we replace Kurumba with other Nilgiri group such as Toda, Kota.

## Results and Discussion

### Maternal diversity and composition

In order to first understand mitochondrial diversity and haplogroup composition, we thoroughly investigated haplogroup composition among nilgiri tribes and its relation to the neighboring populations using 274 complete mtDNA including Toda (N=20), Kota (N=8) and Kurumba (N=109). Generally kurumba comprise many clans (Jenu and Betta Kurumba), but here we took them as a single clan for simplicity. In general three populations posses founder like haplogroup composition like Toda (R5a2b), Kota (M3a1+204), Kurumba (C4a2c1, M2b3a, M36d1, M3c1b1a) (Supplimentary Table 1). The demographic analysis using Bayesian skyline plot indicates that three populations generally possess characteristics of the South Asian population (Welikala et al., 2024; 2026; Desai et al., 2026a; Desai et al., 2026b), where population size (Ne) seem to increase around 40 Kya and remain continue (Supplementary Figure 1ABC). We also noted the decline of population after 5 kya, which might be due to change in lifestyle towards endogamy as indicated by Multiple founder events in different haplogroups in these three groups (Supplementary Figure 2-6). Our broad scale analysis of haplogroup lineage indicates that, generally, Nilgiri groups are closer to neighbour tribes such as Ural Kurumban, and Thogataveera. Importantly, many clades of Toda and Kurumba are shared with groups from Sri Lanka, specifically Indigenous Adivasi (Vedda), which also point towards Nilgiri tribe connection to Sri Lanka (Welikala et al., 2024).

However, one fascinating lineage C4a2c1 found in Kurumba population, which has not been detected in any South Indian population except Kurumba. Generally C4 haplogroup is more restricted to Northeast Asia and Siberia, However, one of its lineages is found only in the Nilgiri tribe, Kurumba. We investigated this lineage thoroughly incorporating aDNA and modern, which indicate that within C4a2a diversified early and found in modern and ancient populations from Russia, Hungary and Mongolia starting from 4800 YBP (Supplementary Table 1; Supplementary figure 6). whereas C4a2b is found in populations of South East Asia and Northeast part of India such as Wanchoo and Sonowal Kachari, this sharing is expected due to Cultural connection associated with Tibeto-Burman groups. The C4a2c bifurcated into two branch around 12 Kya, C4a2c2 lineage predominantly found in Ladakh regions of India, whereas C4a2c1 found in ancient group of Avar from Hungary, Austria, and mostly in ancient population of Xinjiang from China (Jeong et al., 2018; Kumar et al., 2022; Wang et al., 2025). The same branch diverged and entered India around early-mid Holocene (7 Kya), and diversified within India from 5 kya (Supplementary figure 6). It is possible that bearer of this lineage was living North part of Himalayan such Mongolia, one branch migrated towards the Hungary and also remained presented in Xinjiang from China, and specifically C4a2c2 present in Ladakh of India, hence it is plausible that they might have migrated from North of Himalaya to the South of India (Supplementary figure 6).

### Population Structure Among Nilgiri Groups

Our PCA (Principal Component Analysis) using a global population dataset shows that all three Nilgiri groups occupy the broader South Indian population space (Mishra et al., 2024), while exhibiting substantial within-group variation. The Toda form a relatively distinct cluster along the principal axis, with a slight shift away from the East Asian side of the PCA space compared with other South Indian populations (Fig 1). Kurumba and Kota are each further subdivided into two genetic clusters, designated as Kurumba1/Kurumba2 and Kota1/Kota2, respectively according to PCA structure. Kurumba1 is positioned closer to the Toda and towards the Indo-European-related axis, whereas Kurumba2 shifts towards the southern Dravidian-associated axis and clusters more closely with Kota1. Interestingly, Kota2 shows a distinct position towards populations associated with the Austroasiatic groups, suggesting additional genetic affinity that differentiates it from the other Nilgiri groups (Fig 1).

**Fig. 1.**
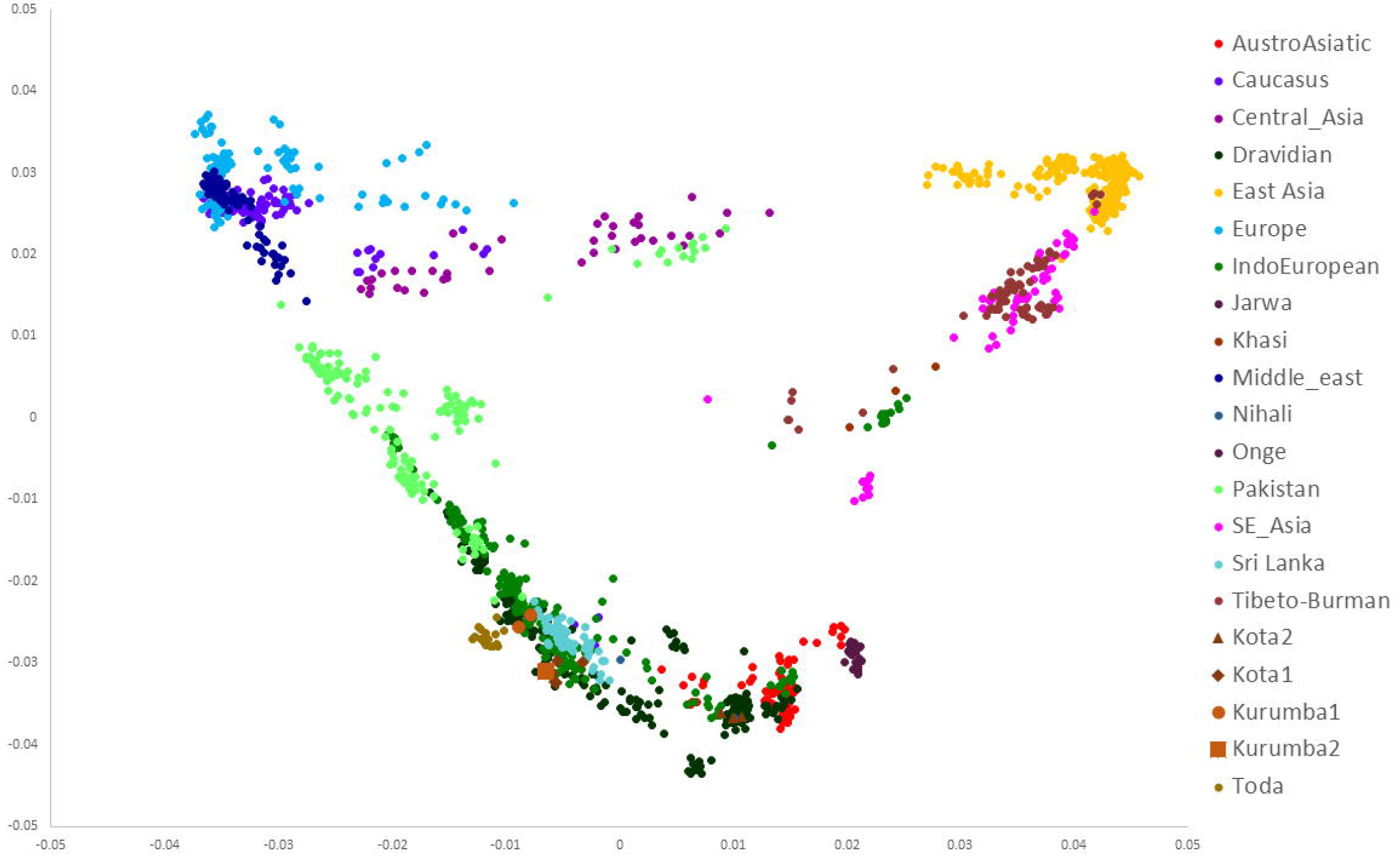
Principal component analysis (PCA) of global populations, showing the genetic position of the Nilgiri populations, their internal substructure, and their relationships with other worldwide populations.

To investigate the broad-scale population admixture history, we performed ADMIXTURE analysis across K = 2–15, with K = 11 selected based on cross-validation error (Fig 2). Overall, the ancestry components recovered at K=11 broadly corroborate previous studies, with Indian populations showing varying proportions of Indigenous South Asian (AASI-related; dark green), Iranian farmer-related (dark blue), and Indo-European-associated (light blue) ancestry, while Austroasiatic and Tibeto-Burman populations show additional East Asian-related ancestry. Within the Nilgiri groups, Toda develops a distinct ancestry profile from K=5 onwards, consistent with its differentiated position in the PCA and potentially reflecting substantial genetic drift and isolation (Fig 2, Supplementary Figure 7-8). Kota1 and Kurumba2 broadly retain ancestry profiles comparable to other regional populations, whereas Kota2 shows an elevated Austroasiatic-associated component. Interestingly, Kurumba1 lacks the teal component almost entirely, in contrast to Kurumba2 and Kota1. This teal component is particularly notable because it is observed mainly in Kurumba2, Kota1, KayaDora1, Gaud1, and a single Mahar individual, and is also enriched in the Vedda population of Sri Lanka, suggesting a potentially shared ancestry component (Fig 2).

**Fig. 2.**
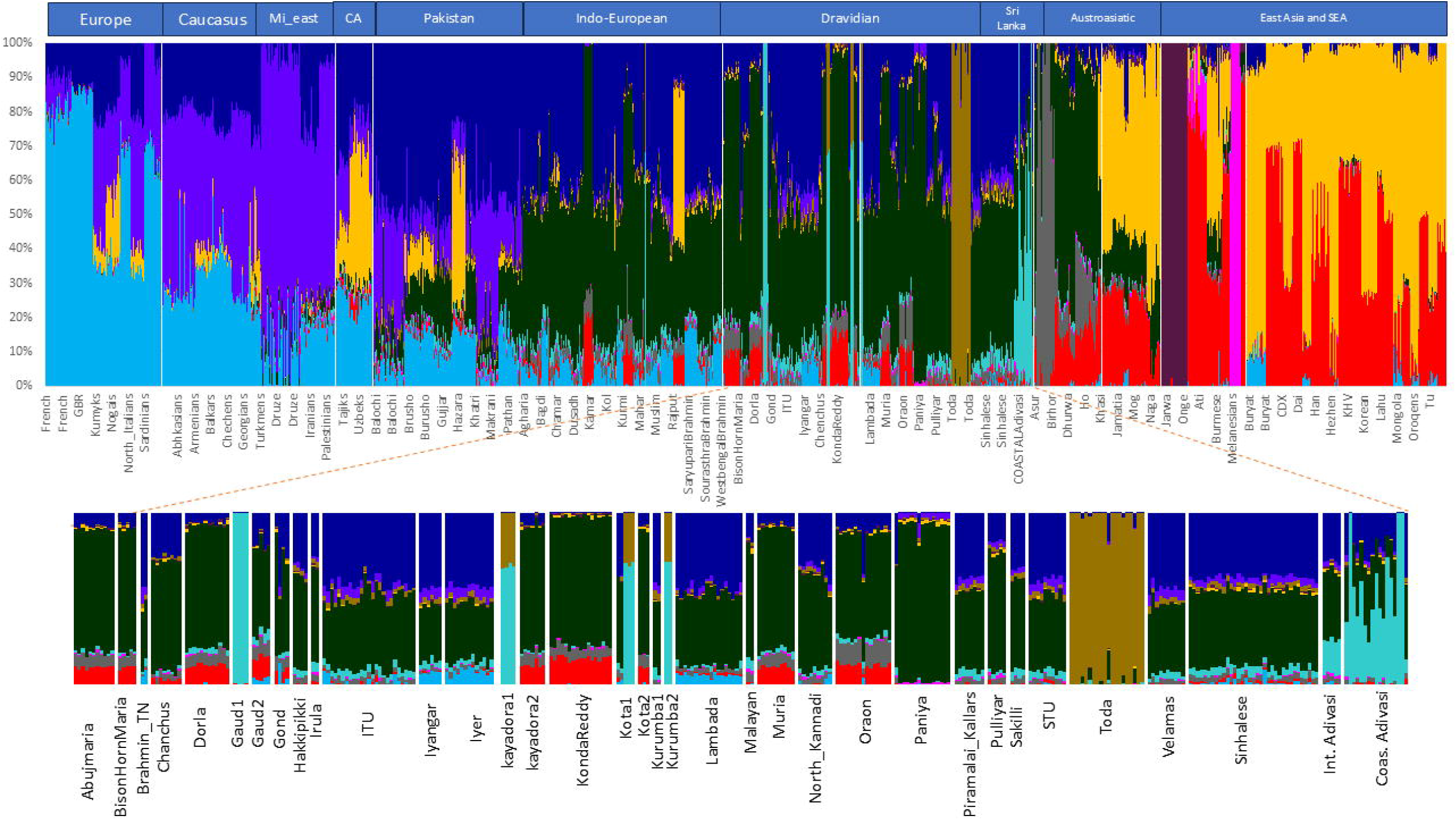
ADMIXTURE analysis at *K* = 11 showing the ancestry composition of the Nilgiri populations and their affinities with populations from South Asia and other geographic regions.

Besides this, we used ASCEND (Tournebize et al., 2022) as we wanted to understand the demographic history of Toda and Kota1, as Kota2 and both Kurumba groups were having sample size below 5, Toda seem to suffer heavy founder event with intensity of 11.8%, with generation at 20, whereas kota1 suffered founder intensity of 4.9% with 25 generation (Supplementary figure 9 and 10). Toda founder event is the same as onge, but with slightly lower intensity (Tournebize et al., 2022), this highlights the distinct demography of Toda and corroborates with the result of Admixture (Fig 2).

### Ancestry connections across the southern Indian Ocean rim

The teal component is more enriched in Gaud1, a population from orissa, India (South-East India), however this component is visible in all Dravidian and Indo-European populations, specifically enriched in Nilgiri groups, and also strikingly visible in Mahar from North India, and importantly from Population of Sri Lanka specially Vedda. Vedda is indigenous adivasi of Sri Lanka, recently studying the existence of internal structure among them as Interior Adivasi and Coastal Adivasi (Urban Aragon et al., 2025). This component is more enriched in coastal vedda groups, which further add validation to existing substructure among vedda people (Urban Aragon et al., 2025). Our outgroup F3 (Paniya, X, YRI) vs F3 (Toda, X, YRI) reveal teal component bearing group get separated and get drifted towards the Toda (Representative of Nilgiri Groups), specially Srilanka groups CoastalAdivasi and Interior Adivasi, which also posses teal component do not get drifted towards Toda, but slightly (Fig 3), this could because of historical gene flow involving Nilgiri populations associated with Teal components. This is also corroborated by fineSTRUCTURE Analysis, where coastal and Interior Vedda occupy different clades while both remain genetically similar to the Sinhalese from Sri Lanka (Fig 4).

**Fig. 3.**
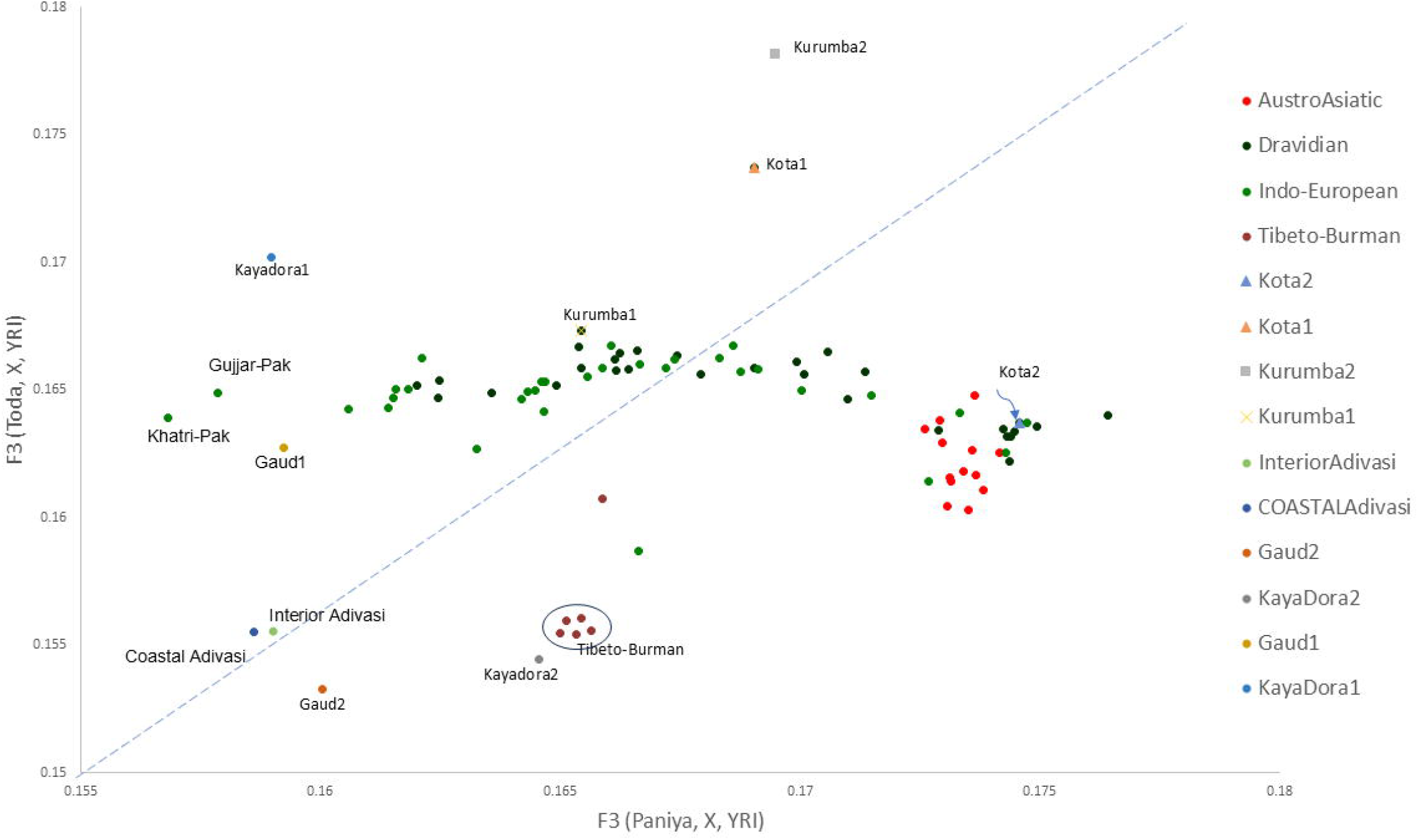
Outgroup *f*3-statistics comparing Paniya and Toda as test populations against a common set of reference populations, illustrating the stratification of Nilgiri and other South Asian groups carrying the teal-associated ancestry component.

**Fig. 4.**
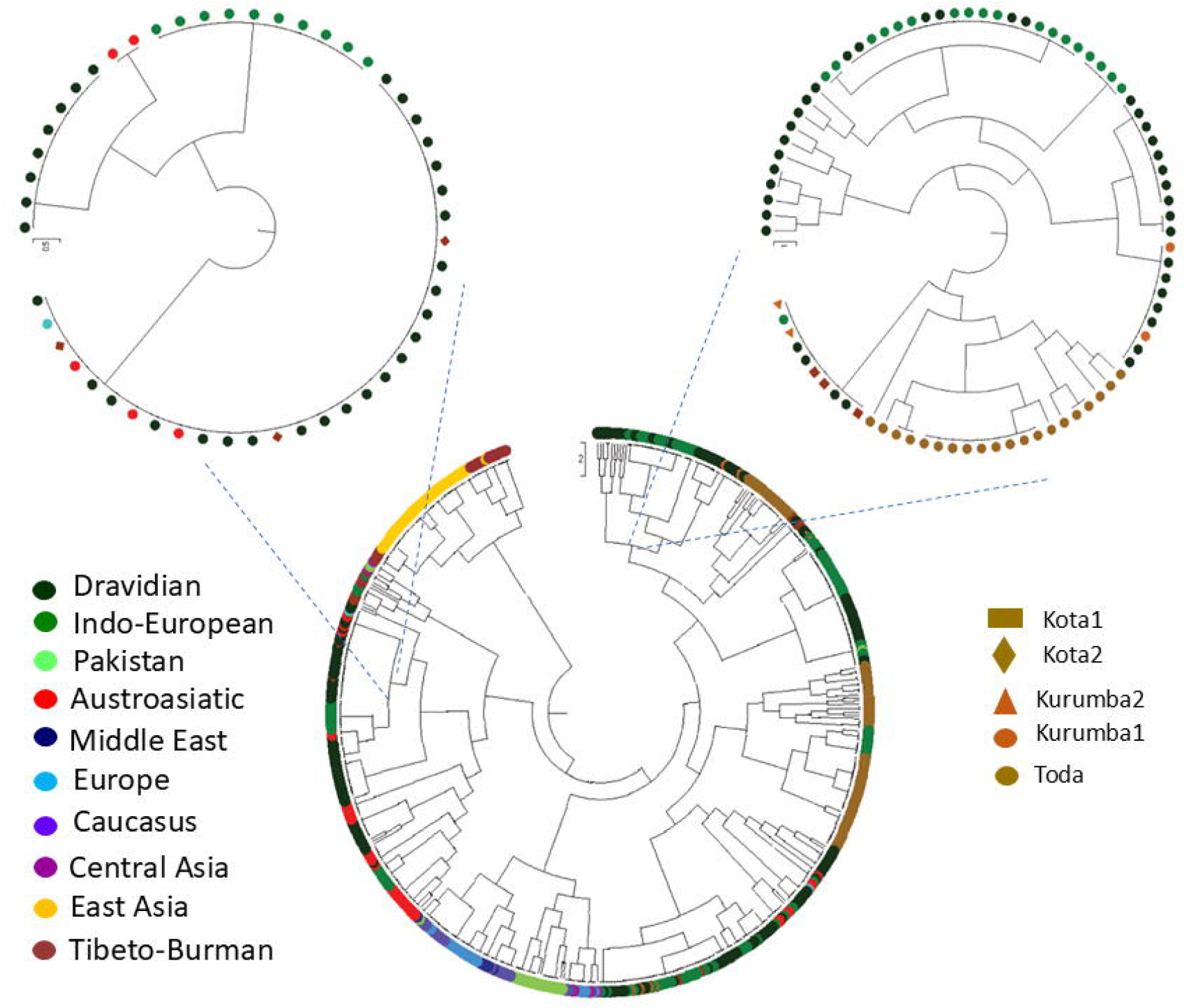
Haplotype-based population structure inferred using fineSTRUCTURE. The fineSTRUCTURE tree reveals internal substructure among the Nilgiri populations and their haplotype-sharing relationships with other populations.

### Nilgiri tribes Affinity to other Populations

Outgroup f3-statistics revealed heterogeneous patterns of shared genetic drift among the South Indian populations. Our Outgroup F3 statistics (Nilgiri groups, X, YRI) revealed that all Nilgiri generally show shared drift with each other and its neighbour Populations (Supplementary Table 1-6). Importantly, Kurumba1 shows high drift with Kurumba2, but Kurumba2 shows more drift to the neighbouring population more, Interestingly with some Indo-European groups such Mahar and Harijan from North India. Specifically Kota2 groups which also show high affinity to the AustroAsiatic in PCA and Admixture, it also shares more alleles to the AustroAsiatic groups (Supplementary Table 4).

In order to investigate the possible North Asia or Siberian ancestry, we performed D statistics in a manner to differentiate Nilgiri groups from others (YRI,Han,X,Kurumba1), however in all tests, results are negative indicating no gene flow or affinity towards Kurumba1 (Supplementary Table 7). Even our test with aDNA sample from different periods of East Asia does not produce any extra affinity. However, we could not see any major difference in these different nilgiri groups to ancient populations. Formal D-statistics revealed significantly negative values across all tested East and Northeast Asian populations in the comparison D(YRI,X;Paniya,Kurumba1) (Z=−6.20 to −9.54) (Supplementary Table 7). This indicates significantly greater allele sharing between Paniya and these East/Northeast Asian populations than between Kurumba1 and the corresponding populations. The consistent pattern across Buryat, Dai, Daur, Han, Hezhen, Japanese, Korean, Mongolian, Naxi and Oroqen suggests that Paniya possesses a stronger affinity to East/Northeast Asian-related ancestry than Kurumba1. This extra affinity could be high due to AASI component in Paniya, as AASI lineage is more related to North Asia lineage, Hence not detecting close affinity to any North east groups to Nilgiri, indicate the dilution of these ancestry among the Nilgiri tribe.

The Nilgiri populations-Kota, Kurumba and Toda-show broadly comparable affinities with the northern Eurasian outgroups, with relatively elevated f3 values for Western Steppe-related populations (WSHG/EEHG, Yamnaya, Srubnaya, Andronovo and Sintashta (Supplementary Table 8). Among them, Toda and Kurumba generally exhibit slightly higher affinity to Western Steppe-related groups than Kota, although the differences are modest. In contrast, Paniya displays a distinct pattern, with comparatively lower affinity to Western Steppe groups but higher shared drift with several Amur and Baikal populations. Overall, the Nilgiri populations do not show a strong, population-specific affinity to any single northern Eurasian source, whereas their relatively elevated affinity to Western Steppe-related groups is expected.

### Ancestry Streams and Genetic Substructure among Nilgiri Populations

As our different combination of D and F3 does not support the extra North east ancestry in autosomal, to assess the number of independent ancestry streams underlying the genetic structure of the Nilgiri populations, we performed qpWave analyses using YRI, WEHG, EEHG, Iran_GanjDareh_N, Anatolia_N, WSHG, ESHG, and Dai as right populations (Table 1). When the genetically defined Kurumba and Kota clusters were analysed in combination with Toda, all four combinations rejected the rank-0 model (P < 10^−33^) but did not reject rank 1 (P = 0.198-0.660), indicating that two independent ancestry streams were sufficient to explain each three-population configuration. In contrast, when all five genetic clusters (Kurumba1, Kurumba2, Kota1, Kota2, and Toda) were analysed simultaneously, both rank 0 and rank 1 were rejected (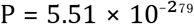 and 0.0096, respectively), whereas rank 2 was not rejected (P = 0.622), indicating a minimum of three independent ancestry streams among the five groups (Table 1). To determine whether the two-stream configuration was specific to the Nilgiri populations, we performed equivalent analyses using selected neighbouring Indian populations in place of the Kota clusters. Kurumba1 paired with Paniya, Mahar, Irula, and Hakkipikki consistently rejected rank 0 while retaining rank 1 (P = 0.600-0.917), indicating that these populations could likewise be explained by two independent ancestry streams. Thus, the two-stream configuration observed among the Nilgiri populations is not unique to them and is also observed among several neighbouring South Asian populations. However, the requirement for three ancestry streams when the internal Kurumba and Kota subdivisions are considered simultaneously highlights additional genetic complexity within the Nilgiri groups.

**Table 1.**
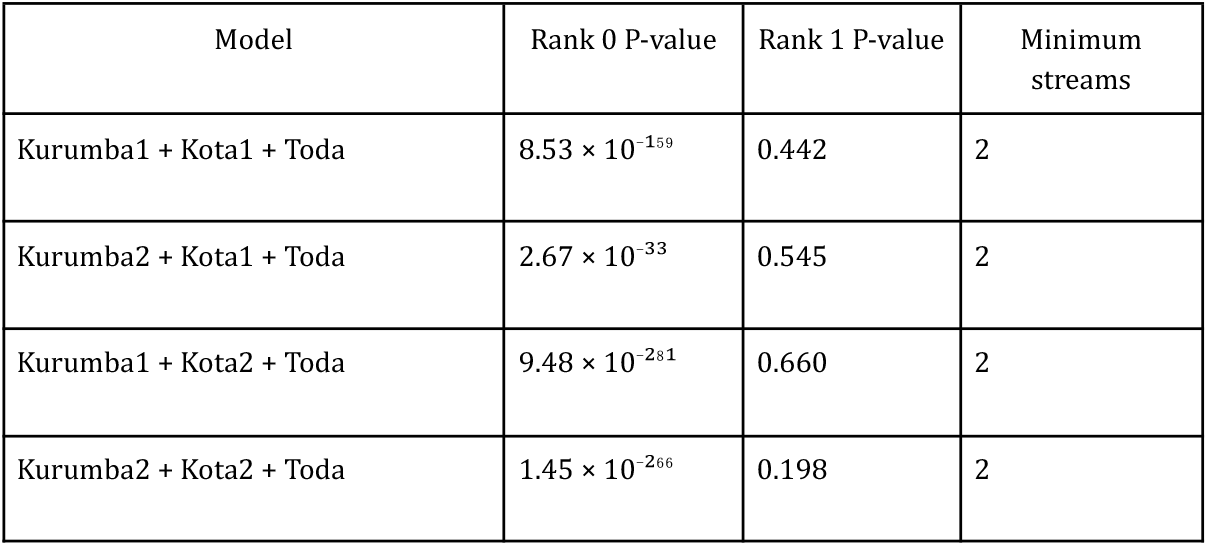
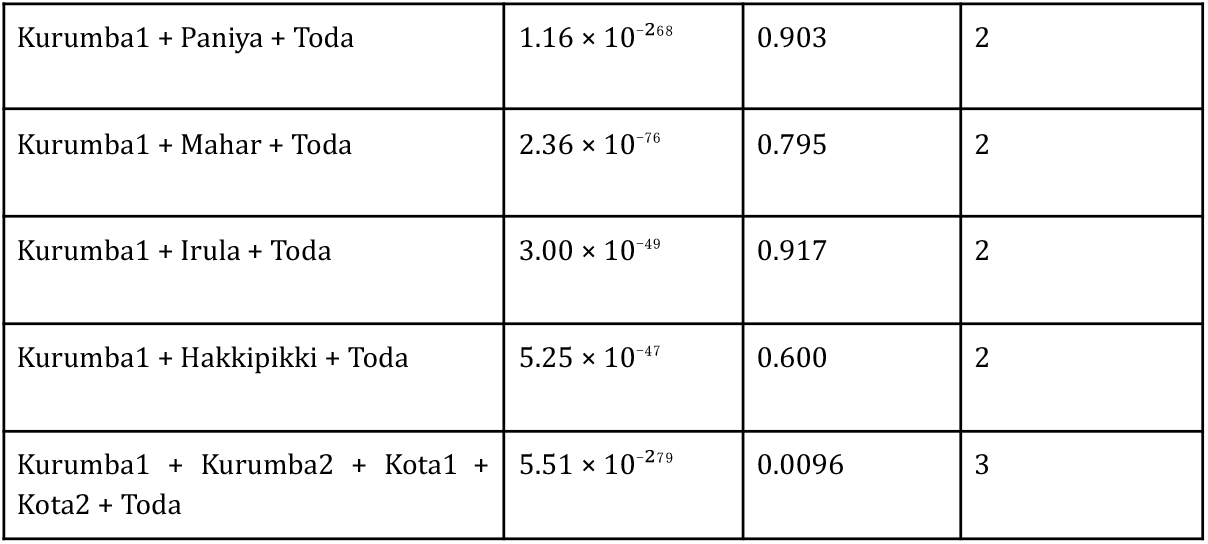
Table showing the different tested qpWave model with respect to different Nilgiri Tribe.

The qpAdm analysis models the selected South Asian populations as mixtures of Onge-, Indus_Periphery_HighCoV, and Western Steppe MLBA-related ancestry. Across the tested groups, the Onge-related component generally contributes the largest proportion (Figure 5), whereas the Indus Periphery-related ancestry forms a substantial second component, particularly in the Nilgiri populations. Kurumba1 and Kurumba2 show broadly similar ancestry profiles, with ∼39-44% Onge-related, ∼45-53% Indus Periphery-related, and ∼8-11% Steppe-related ancestry. In contrast, Toda shows a higher Indus Periphery-related contribution (∼59%) and lower Onge-related ancestry (∼33%), with ∼8% Steppe-related ancestry. Among the neighbouring/regional populations, Paniya and Pulliyar show comparatively high Onge-related proportions (∼73% and ∼64%, respectively) (Figure 5), whereas Hakkipikki, Chanchu, and Lambada show intermediate profiles. The model fits are statistically acceptable for most populations, although several groups show marginal or poor fits (P ≤ 0.05), indicating that the three-source model does not adequately explain all groups. Notably, Kota2 has a slightly negative Steppe coefficient and is therefore displayed as zero in the figure, suggesting that the data do not require a Steppe-related contribution under this model. This also explains that Kota2 groups show extra affinity for AustroAsiatic groups and does not form the same cline like other Nilgiri groups, this also confirms the large sub-structure among different groups of Nilgiri populations.

**Fig. 5.**
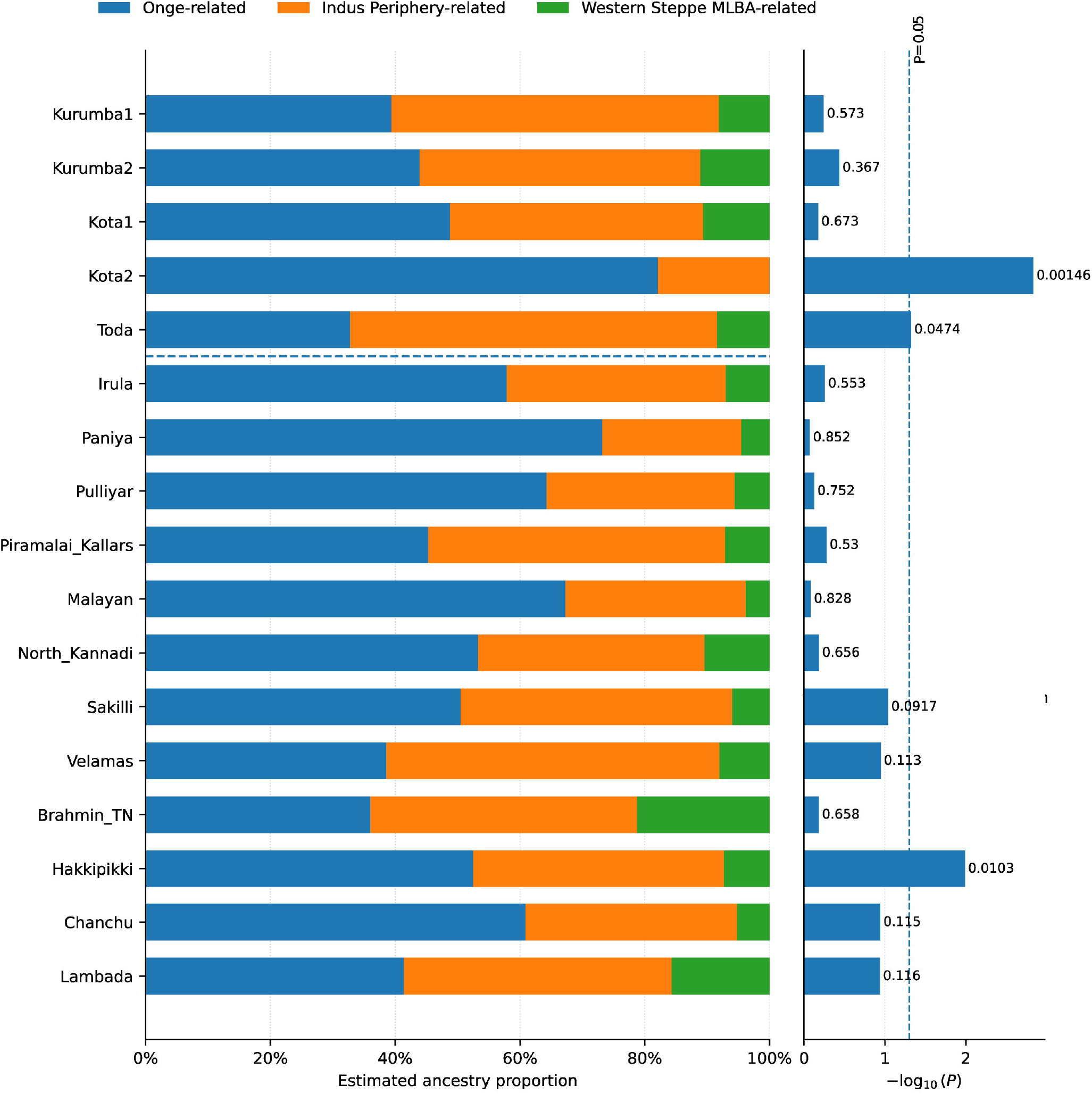
qpAdm-based admixture modelling of the Nilgiri and other South Asian populations using proximal ancient populations as candidate ancestral sources.

## Conclusions

The Nilgir hills of South India hold very important and unique tribes, each complementing in function and co-existing for a long time. However, till now they have been understudied from genetic perspective. This present study reveals the existence of sub-structure among these populations. Specifically striking sub-structure among Kota group, where kota2 show high affinity to the AustroAsiatic groups. The unexpected haplogroup C4a2c presently only in Kurumba, but any Nilgiri does not show the extra affinity to North Asia or Siberian populations, and this haplogroup seems to enter India around the holocene period from North India. This also indicates that maternal signature persisted but nuclear signals seem to dilute after long generation time. The unique teal component shared among the population from South East-South West, North India and Sri lanka population seem to be an ancient connection between these groups, whereas lack of high drift sharing between teal component harbouring groups, and Sri Lanka group, may be lack of recent gene flow. Overall, Nilgiri tribe possesses very unique structure and ancestry among them, but connected to their neighbour more through geography.

## Contributions

SD, RT and GC conceived and designed the study. SD, RT, MKT, AW and GC were involved in performing the analyses. SD carried out the primary data processing and statistical analyses. SD, and GC wrote the original draft of the manuscript. All authors contributed to data interpretation, critically revised the manuscript for intellectual content, and approved the final version.

## Acknowledgments

S.D. is supported by the Council of Scientific and Industrial Research-Senior Research Fellowship (SRF-CSIR). G.C. is supported by Indian Council of Medical Research ad hoc grants (2021-6389 and 2021-11289) and Institute of Eminence, Banaras Hindu University (6031). The support and resources provided by the PARAM Shivay Facility under the National Supercomputing Mission, Government of India at the Indian Institute of Technology, Varanasi are gratefully acknowledged.

