## Supplemental Figure 1-10 for "Holocene North Asian gene flow and cryptic substructure in the genome history of Nilgiri tribes"

*Supplementary Figure 1A. Bayesian skyline plot of Kota groups using complete mtDNA sequences.*

*Supplementary Figure 1B. Bayesian skyline plot of Kurumba groups using complete mtDNA sequences.*

*Supplementary Figure 1C. Bayesian skyline plot of Toda groups using complete mtDNA sequences.*

*Supplementary Figure 2. The bayesian tree analysis of R5 Haplogroup of Toda population and its affinity to neighbour groups.*

*Supplementary Figure 3. The Bayesian Tree analysis of the M36 haplogroup of Kurumba, which shows founder events.*

*Supplementary Figure 4. The Bayesian Tree analysis of the M35 haplogroup of Kurumba, which shows founder events.*

*Supplementary Figure 5. The Bayesian Tree analysis of the M2 haplogroup of Kurumba, which shows founder events.*

*Supplementary Figure 6. The Bayesian Tree analysis of the C4a2b haplogroup of Kurumba, indicating holocene migration to India.*

*Supplementary Figure 7. Plot showing the admixture from K=2 to K=8, indicating how population structure is changing with increasing number of components and admixture between different populations.*

*Supplementary Figure 8. Plot showing the admixture from K=9 to K=14, indicating how population structure is changing with increasing number of components and admixture between different populations.*

*Supplementary Figure 9. Founder event time and intensity of Kota1 groups estimated using ASCEND.*

*Supplementary Figure 10. Founder event time and intensity of Toda groups estimated using ASCEND.*

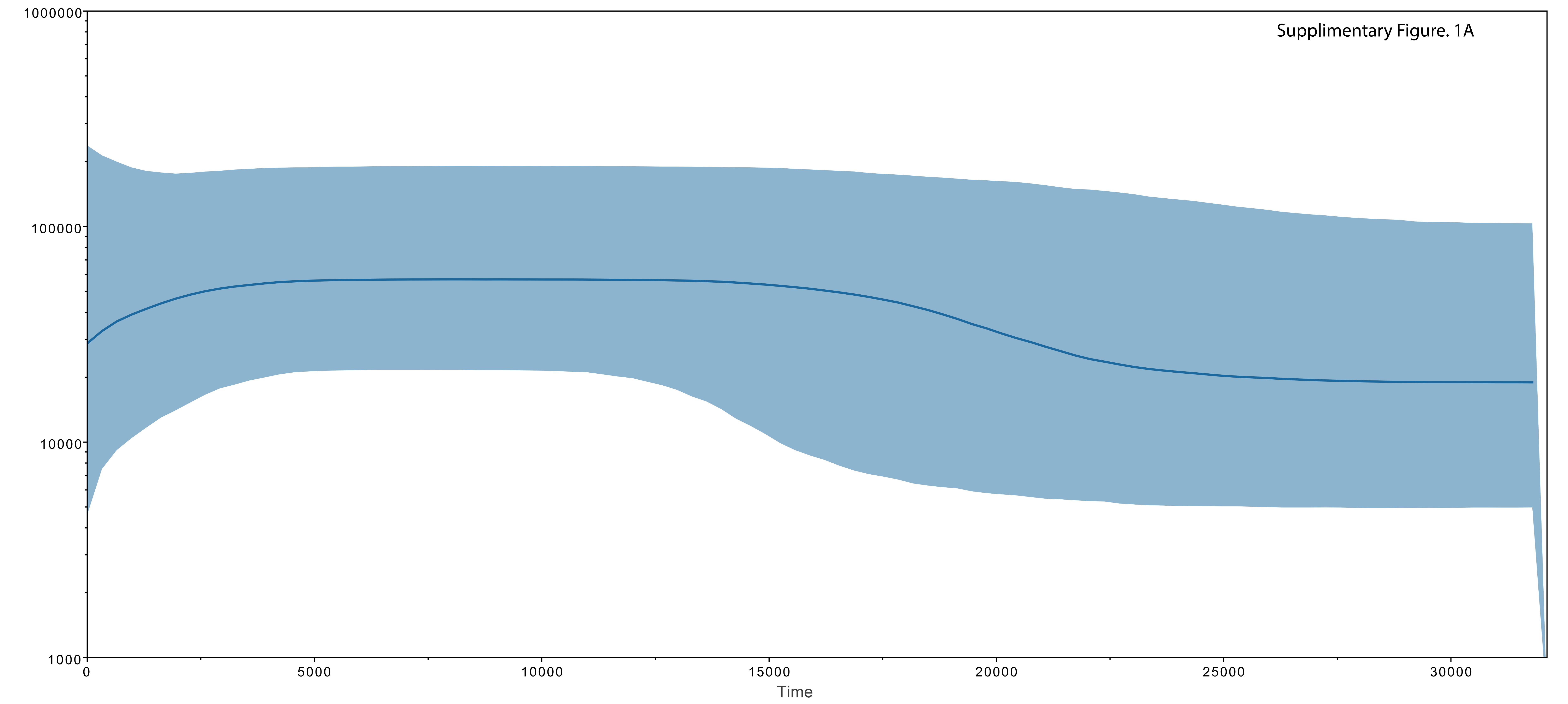
Supplimentary Figure. 1A
1000000
100000
10000
1000
0
5000
10000
15000
20000
25000
30000
Time

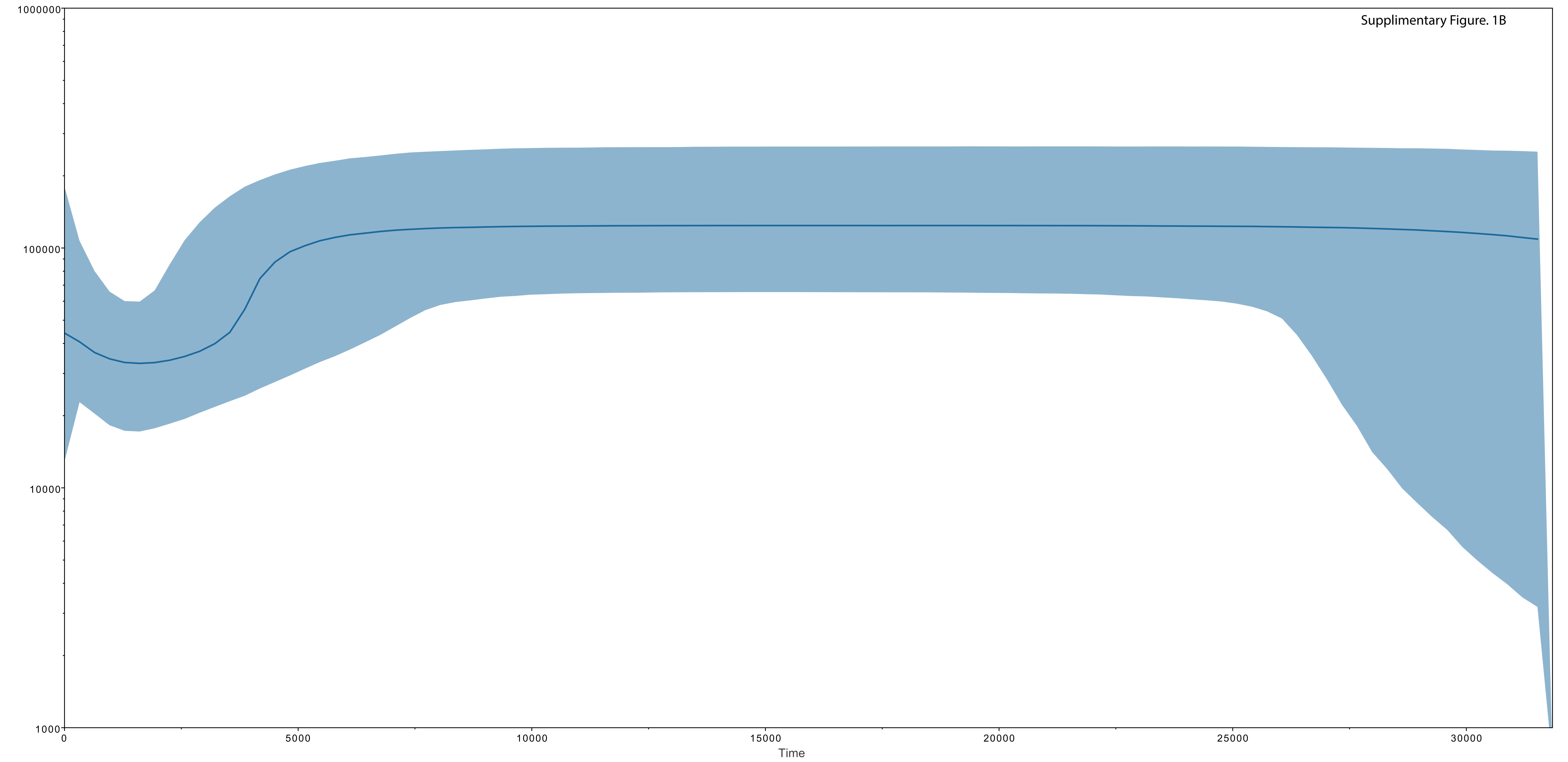
Supplimentary Figure. 1B
1000000
100000
10000
1000
0
5000
10000
15000
20000
25000
30000
Time

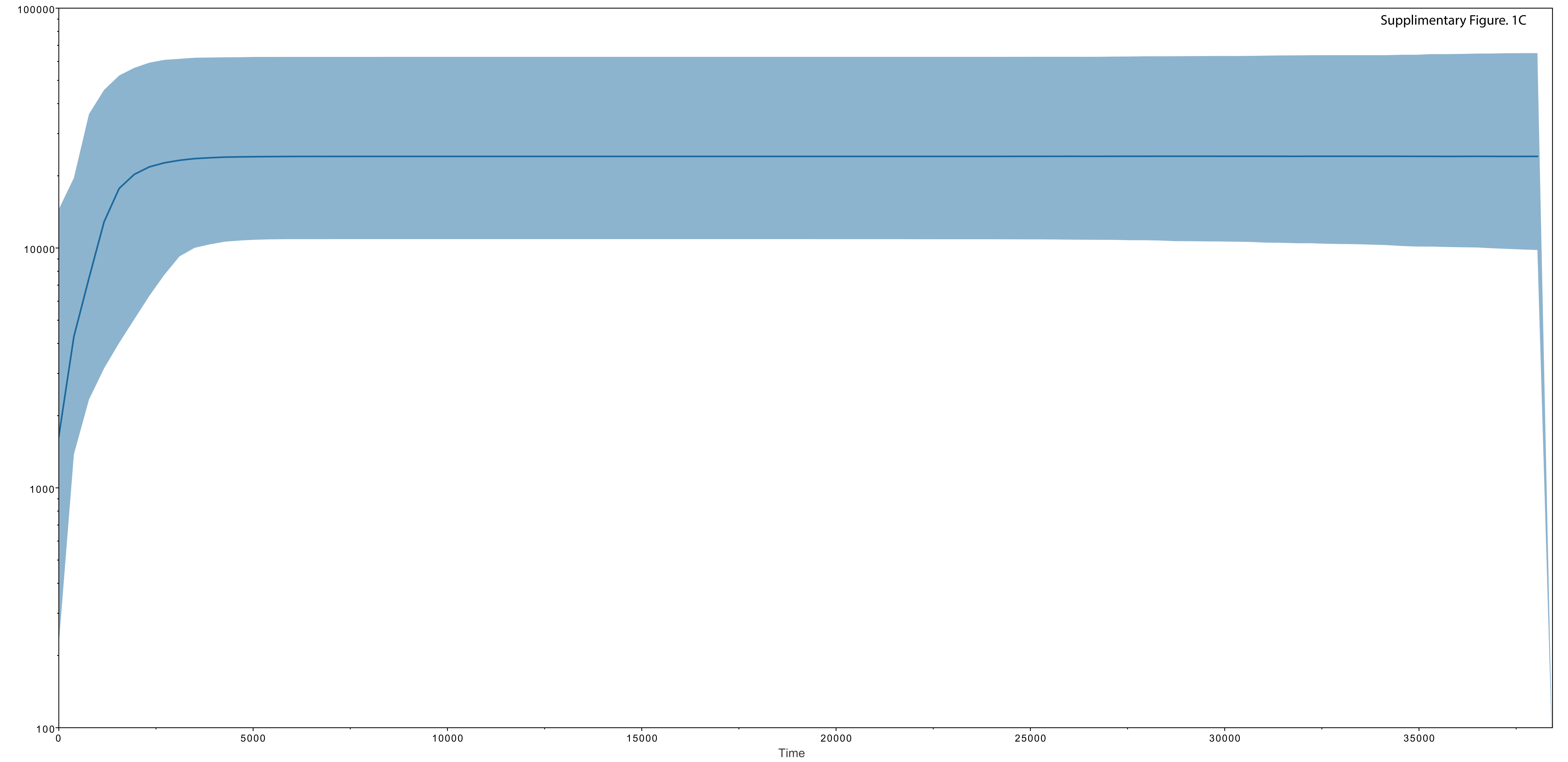
Supplimentary Figure. 1C
100000
10000
1000
100
0
5000
10000
15000
20000
25000
30000
35000
Time

Supplimentary Figure. 2

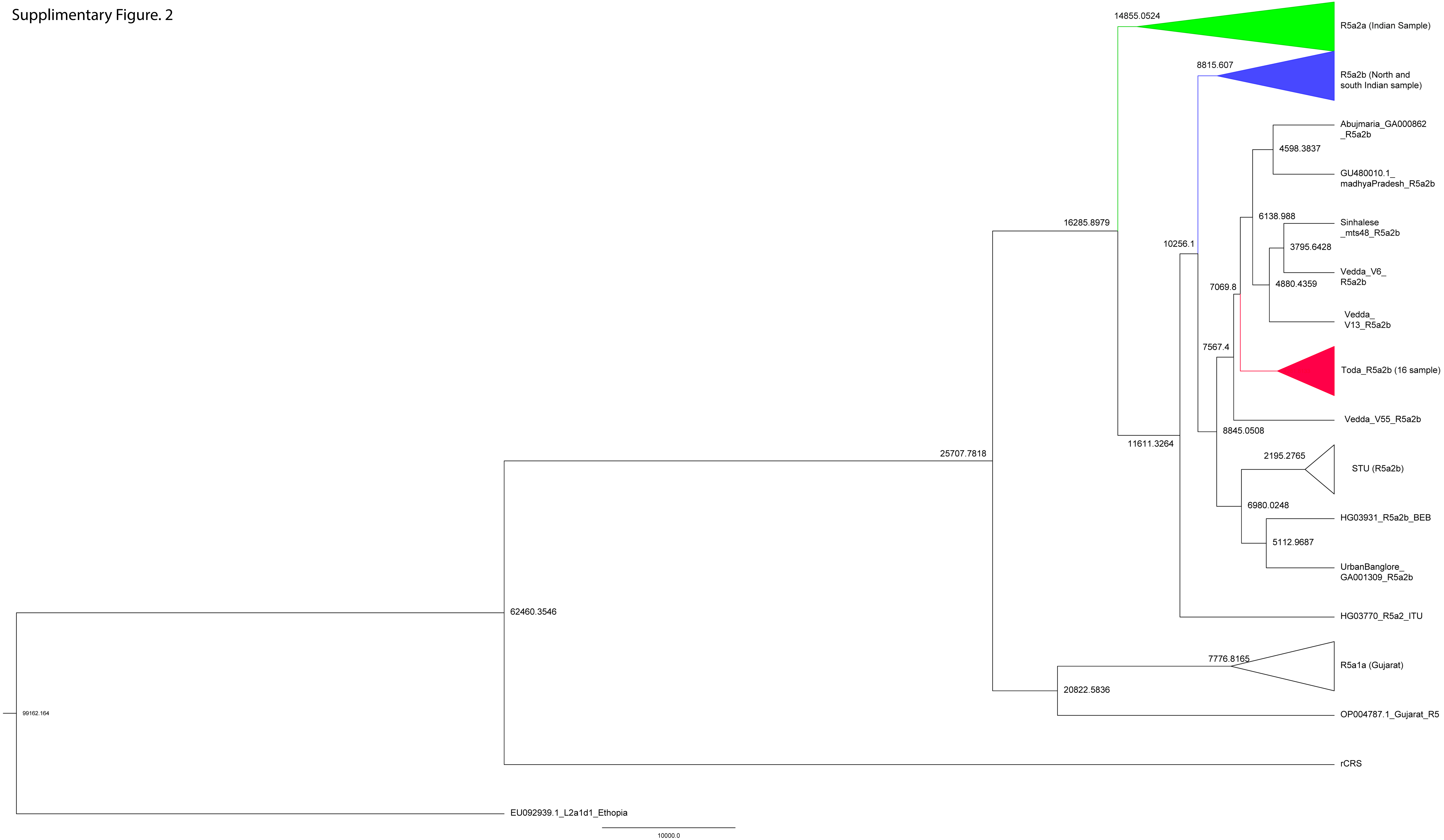

Supplimentary Figure. 3

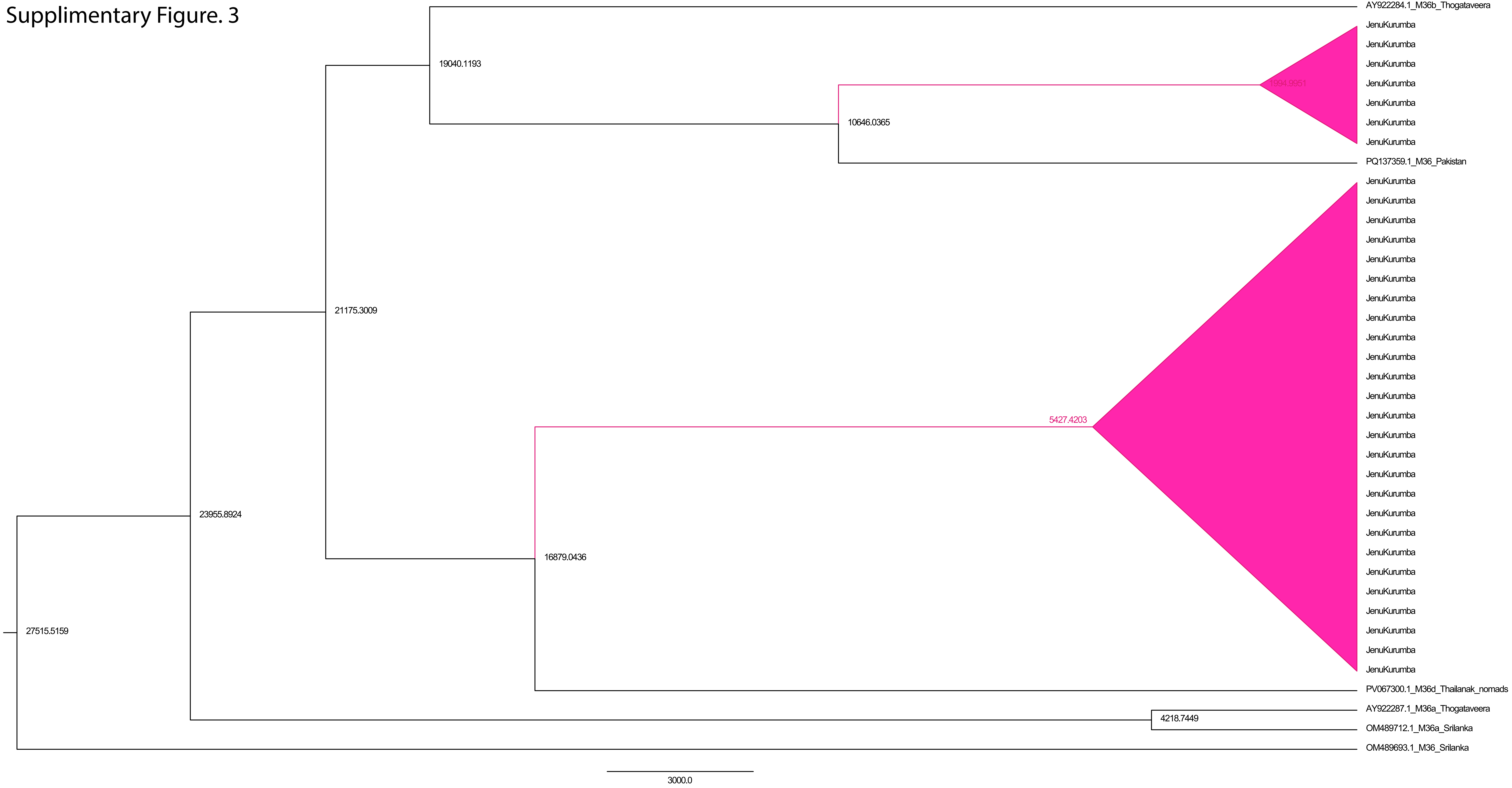

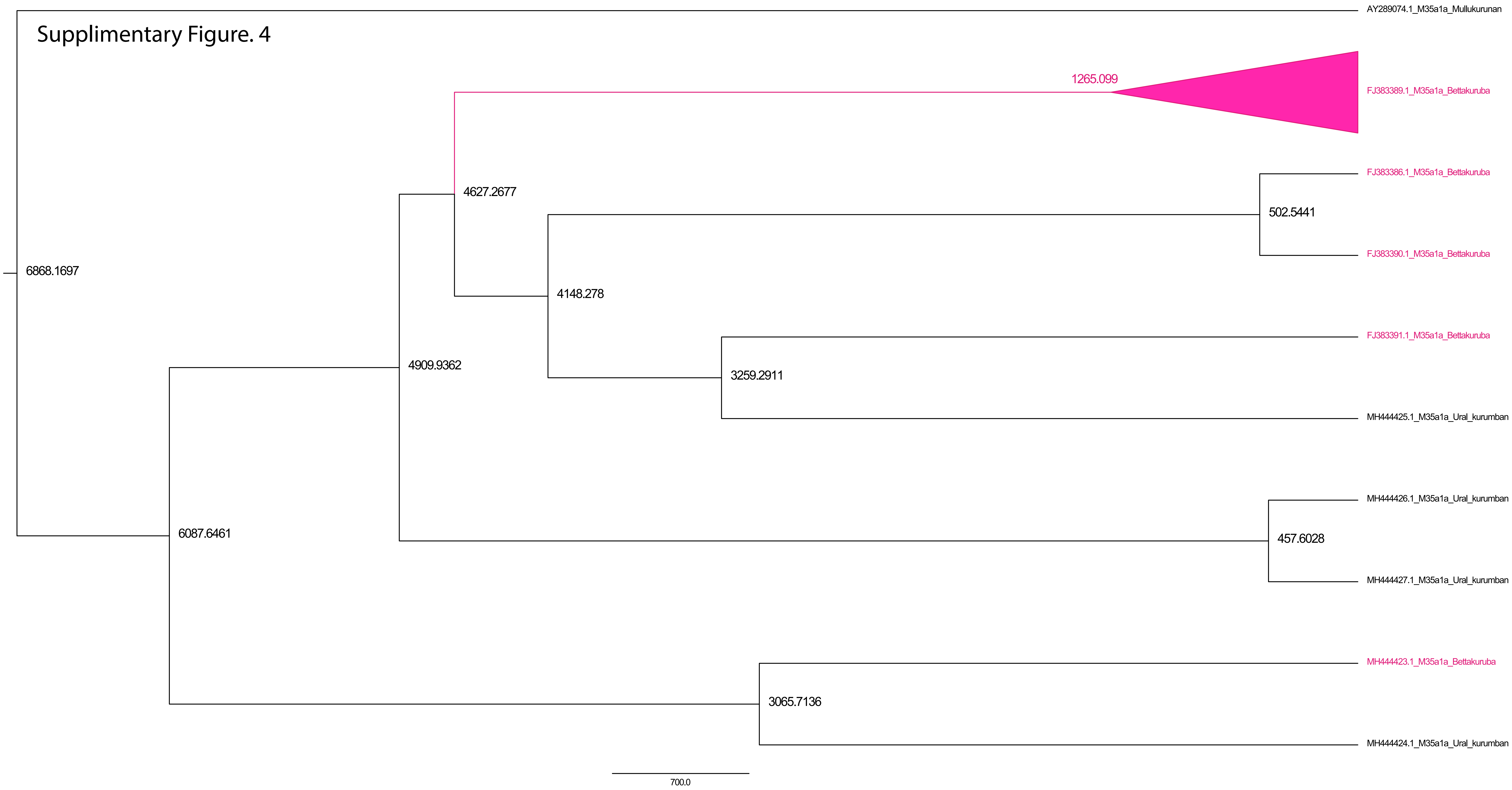

Supplimentary Figure. 4
AY289074.1_M35a1a_Mullukurunan
1265.099
FJ383389.1_M35a1a_Bettakuruba
FJ383386.1_M35a1a_Bettakuruba
4627.2677
502.5441
FJ383390.1_M35a1a_Bettakuruba
6868.1697
4148.278
FJ383391.1_M35a1a_Bettakuruba
4909.9362
3259.2911
MH444425.1_M35a1a_Ural_kurumban
MH444426.1_M35a1a_Ural_kurumban
6087.6461
457.6028
MH444427.1_M35a1a_Ural_kurumban
MH444423.1_M35a1a_Bettakuruba
3065.7136
MH444424.1_M35a1a_Ural_kurumban
700.0

Supplimentary Figure. 5

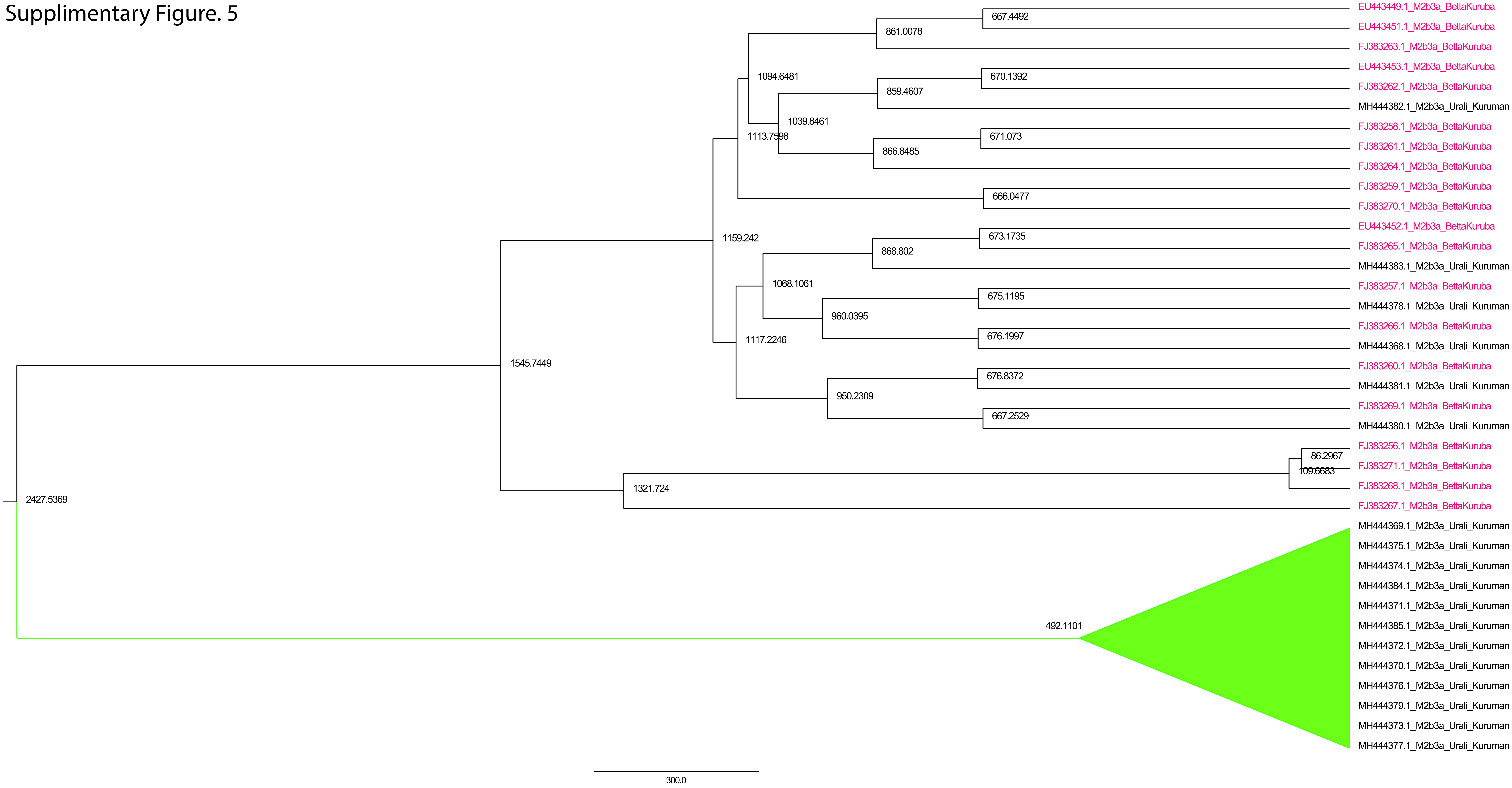

Supplimentary Figure. 6

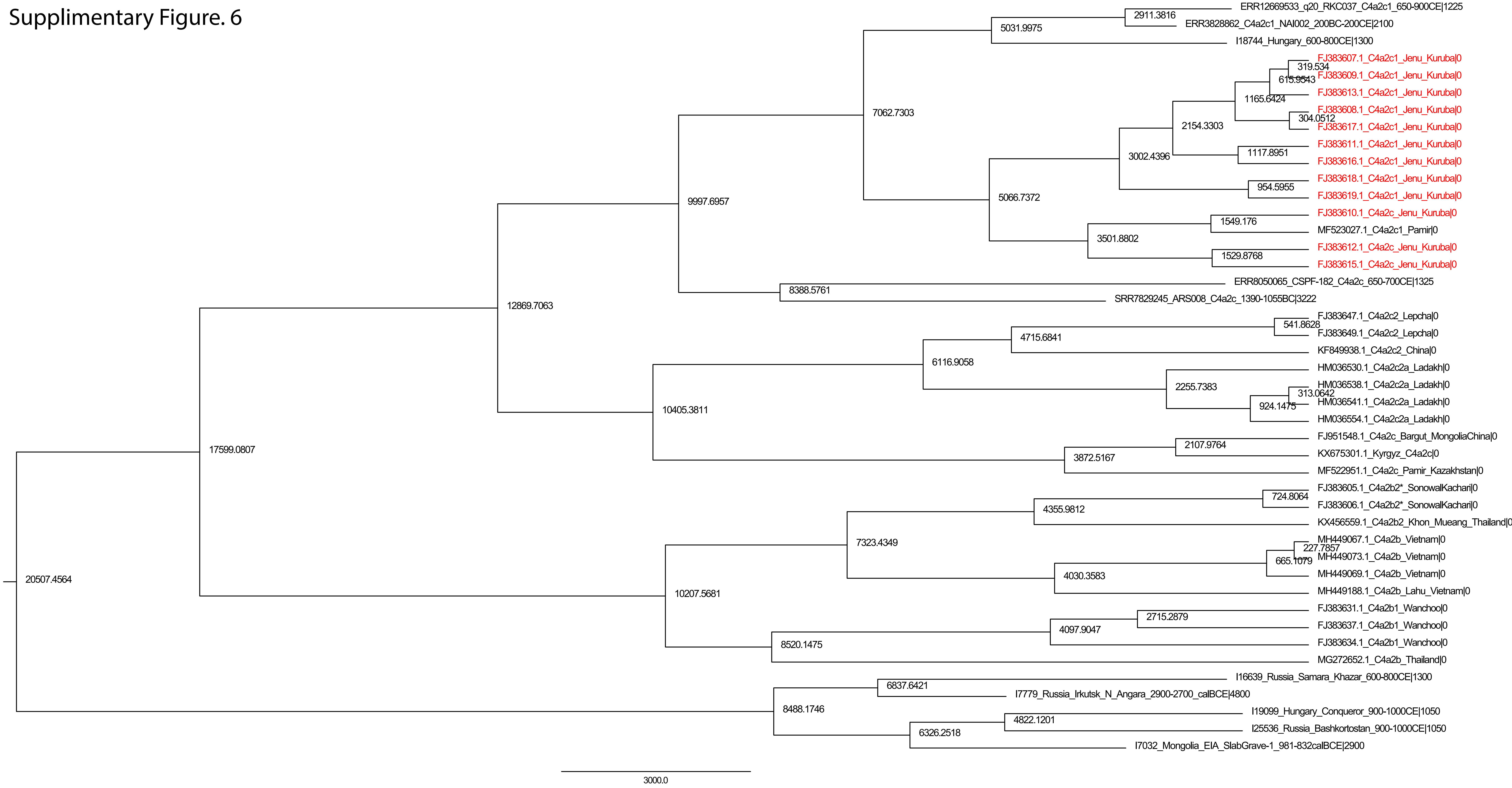

Supplimentary Figure 7

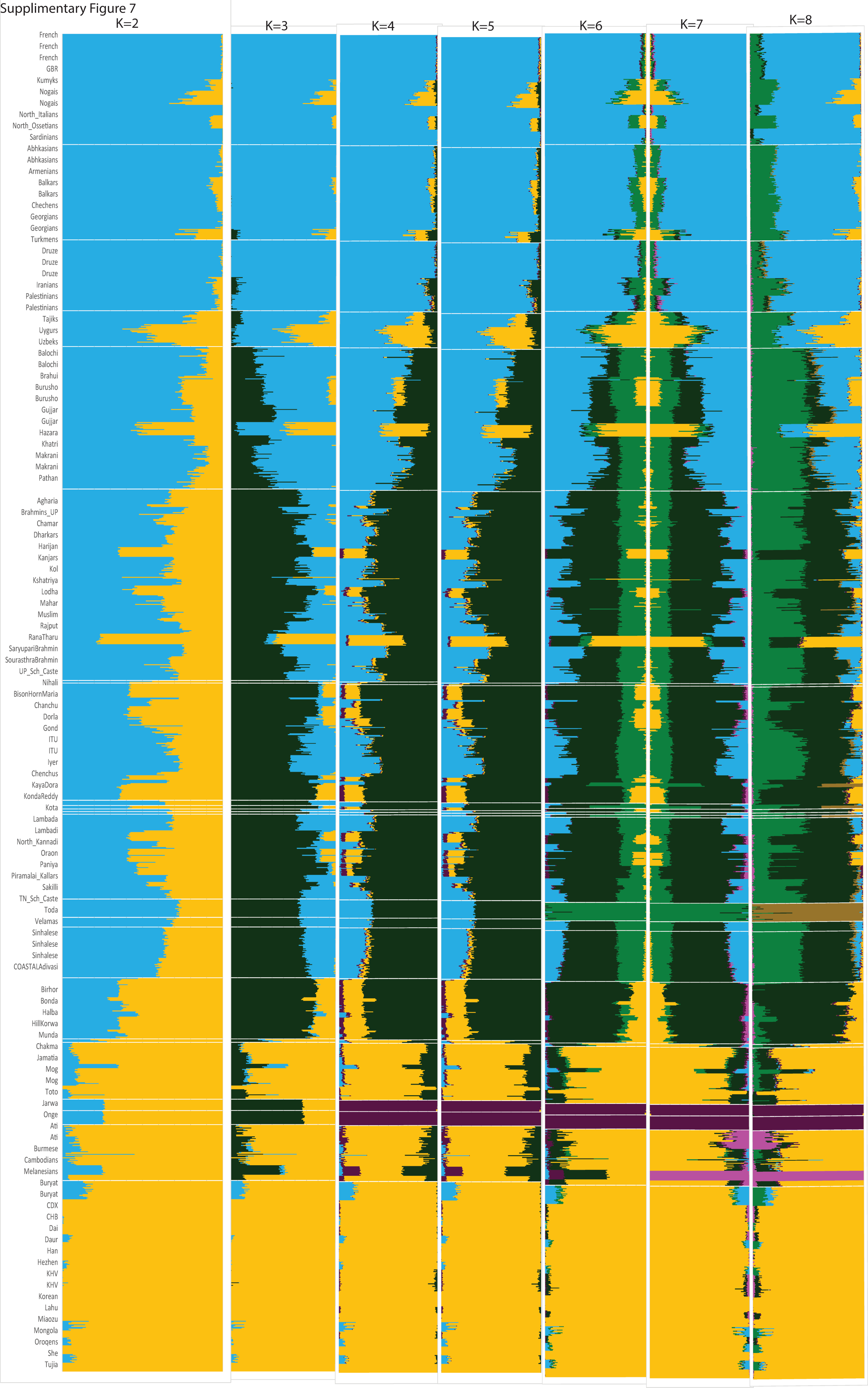

Supplimentary Figure 8

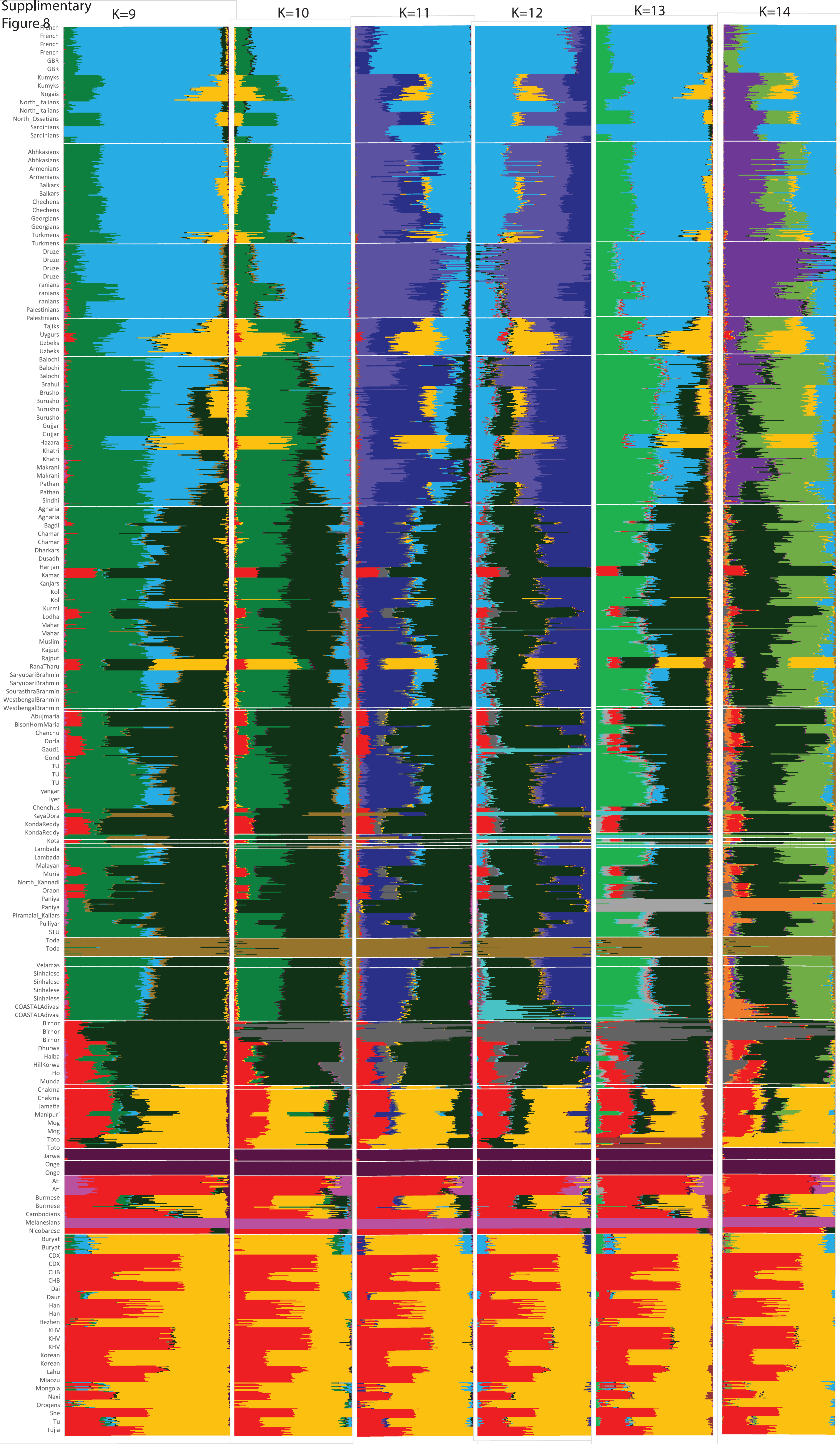

Supplementary Figure 9

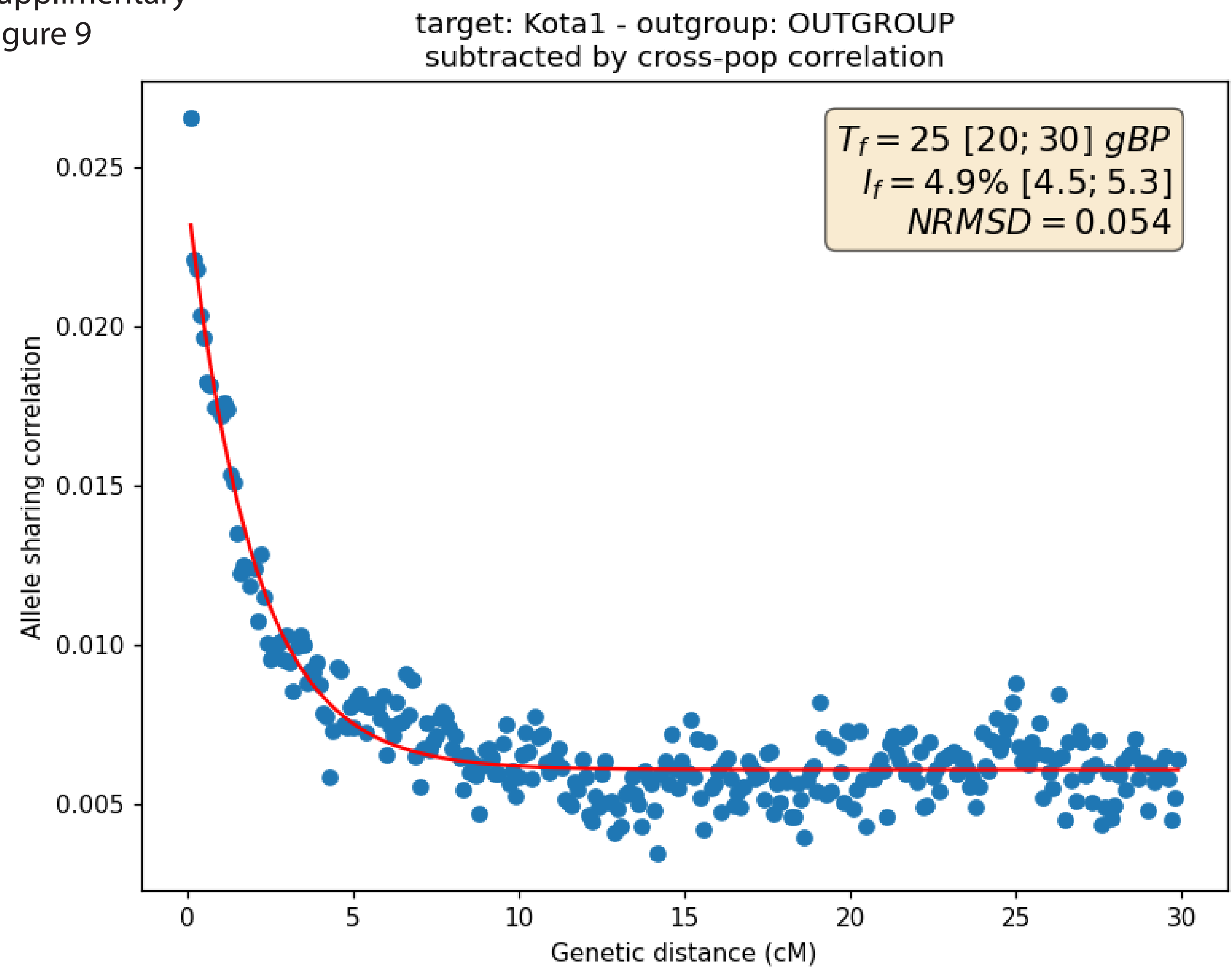


Supplementary Figure 10

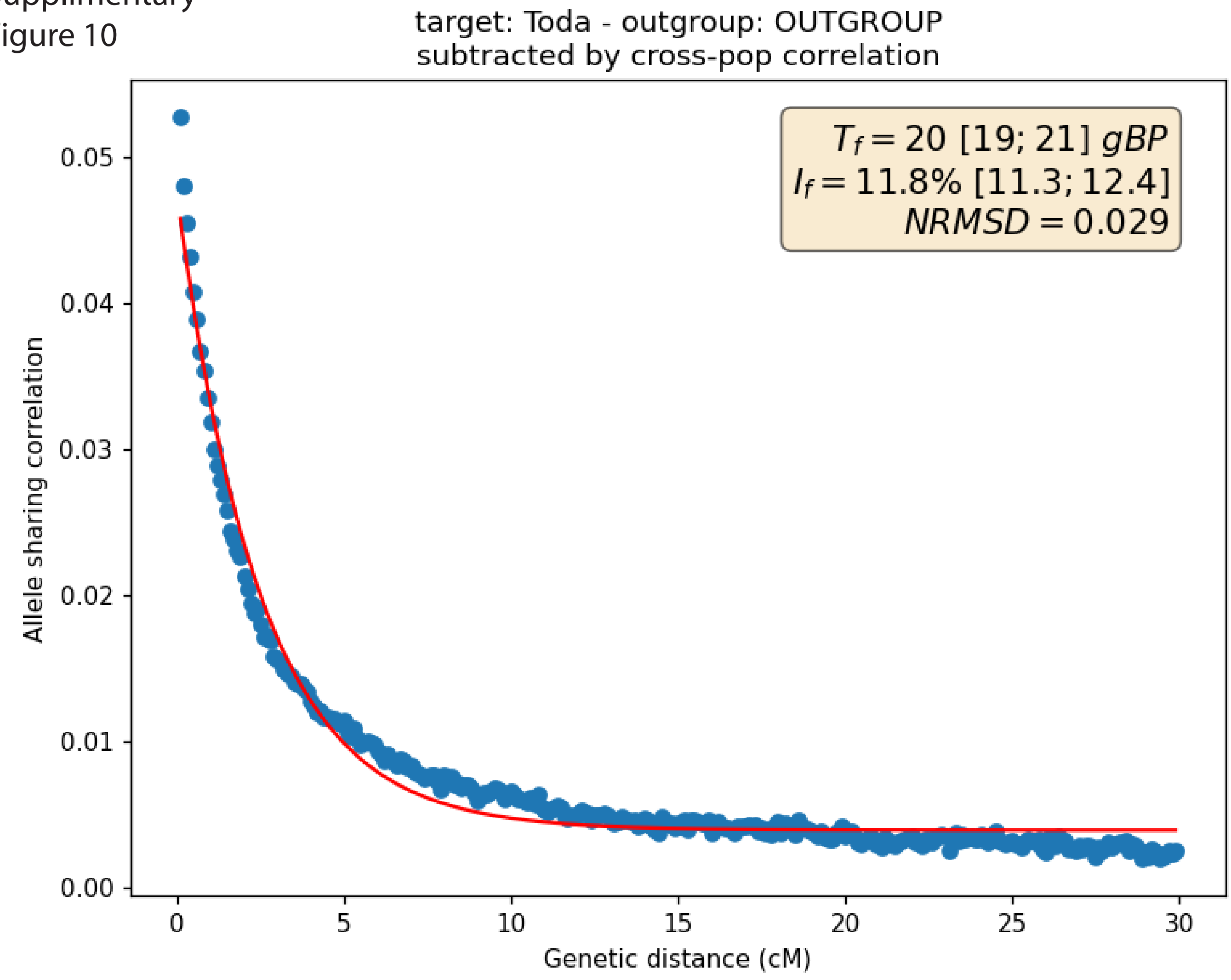
